# Phylogeny Inference Under Time-Decaying Migration and Varying Information Content

**DOI:** 10.1101/2023.11.23.568248

**Authors:** Zaynab Shaik, Nicola G. Bergh, G. Anthony Verboom, Bengt Oxelman

## Abstract

Postspeciation gene flow is widespread across the Tree of Life but is ignored as a cause of gene tree discordance under the standard multispecies coalescent. Where interspecific migration has occurred but is not modelled explicitly, effective population sizes, divergence times and topology can be seriously misestimated. Isolation-with-migration and multispecies coalescent-with-introgression models explicitly model migration but include additional parameters that limit their computational viability with even moderately sized molecular data sets. Here we simulate the evolution of sequences which vary in molecular information content under the coalescent while allowing continuous, tree-wide gene flow/migration between contemporaneous branches, the rate of which decreases with time since divergence. Using simulated sequences, we evaluate the performance of DENIM under rapidly to gradually time-decaying migration and benchmark its performance against the standard MSC method StarBeast3. DENIM consistently outperforms StarBeast3, both in phylogenetic accuracy and computational performance per core. Rapidly decaying migration is associated with improved topology and divergence time estimates under both DENIM and StarBeast3. While species tree estimation accuracy is not improved by increasing the number of loci from 30 to 60 under either method, model convergence is slowed considerably. By contrast, increasing sequence length to 10,000 bp has no clear effect on convergence rates, but shows a tendency towards increased accuracy in DENIM. We apply DENIM and StarBeast3 with a 36-locus empirical bat data set and recover species trees identical in topology to those obtained with 12,931 loci. Our work demonstrates that DENIM can deliver accurate phylogenetic estimates in the presence of both deep coalescence and empirically realistic migration patterns using as few as 30 loci with single-core runtimes of 2-3 days.

---

The scale of genome-wide molecular data and the development of increasingly sophisticated Bayesian models for phylogeny reconstruction using Markov chain Monte Carlo (MCMC) have outpaced the computational resources typically available to molecular phylogeneticists. Fully parameterised Bayesian species tree inference software (e.g., Liu & Pearl 2007; Heled & Drummond 2010; Bryant et al. 2012; Yang & Rannala 2010, 2014; Jones et al. 2015; Yang 2015; Jones 2017a; Ogilvie et al. 2017; Douglas et al. 2022) implement the Multispecies Coalescent (MSC), an extension of the population genetic n-coalescent (Kingman 1982), to model the genealogical relationships among species. The MSC relaxes the assumption that gene trees share a common genealogical history (Ogilvie et al. 2016), allowing them instead to follow a coalescent process wherein the topologies and branch lengths of alternative gene trees vary. This explicit acknowledgement of genealogical discordance arising from incomplete lineage sorting (ILS) has been called a major advance in molecular phylogenetics (Edwards 2009), but critically, the MSC ignores alternative causes of gene tree discordance, including post speciation gene flow, as well as gene duplication and loss with subsequent misinterpretation of paralogues as orthologues (Maddison 1997; Rasmussen & Kellis 2007).

The MSC assumes instantaneous termination of gene flow at the splitting time between two lineages (Rannala & Yang 2003), but empirical evidence from across the Tree of Life provides support for speciation instead being a gradual process during which diverging lineages frequently exchange genes (Nosil 2008; Martin et al. 2013; Liu et al. 2014; Xu & Yang 2016; Nielsen et al. 2017; Burbrink & Gehara 2018; Mao et al. 2018; Morales & Carstens 2018; Thawornwattana et al. 2018; Wu et al. 2018; Pavón-Vázquez et al. 2021; Suvorov et al. 2022; for reviews see Mallet et al. 2016; Taylor & Larson 2019; Edelman & Mallet 2021). Evidence from simulated and empirical data demonstrates that failing to account for migration where it has occurred leads to systematic overestimation of ancestral population sizes, underestimation of speciation times, and, under certain conditions, errors in topology estimation (Leaché et al. 2014a; Long & Kubatko 2018; Müller et al. 2018; Wen and Nakhleh 2018; Elworth et al. 2019, Jiao et al. 2020, Pang & Zhang 2022).

Efforts to model interspecific migration in tree inference have culminated in two extensions of the standard MSC with added migration parameters. The first of these, comprising isolation-with-migration (IM) methods and MSC-with-migration methods (Hey & Nielsen 2004), models migration as a continuous process. Besides effective population size and divergence time estimates in the standard MSC, IM methods estimate a rate of migration per generation *M* in either direction for every pair of contemporaneous branches in the species tree, totalling 2(*n*-1)^2^ migration rate parameters for a tree with *n* tips (Hey 2010; Jones 2019a). A special case of the IM model is the isolation-with-initial-migration model (IIM; Wilkinson-Herbots 2012; Costa & Wilkinson-Herbots 2017) that enables continuous migration at rate *M* from the time of divergence *τ*_0_ until time *τ*_1_ > 0, at which point gene flow between the branches ceases. The second extension of the standard MSC models migration discretely and comprises MSC with introgression (MSCi) and multispecies network coalescent models (MSNM; Rannala et al. 2020). MSCi models are more parameter-rich than IM methods because both the timing *τ* and probabilities (φ or γ) of discrete reticulation events are estimated (Flouri et al. 2020; Rannala et al. 2020). If extinct or unsampled “ghost” lineages are explicitly represented in a network by non-horizontal reticulation edges, the branch lengths of these additional edges also require estimation (Degnan 2018). While both IM and MSCi methods accommodate post speciation migration, IM models are only appropriate where evolutionary relationships can be represented by a bifurcating tree. Where two lineages merge to form one, a network is necessary to represent evolutionary relationships appropriately (Bravo et al. 2019). Allopolyploidy can be represented by a network, or alternatively by a bifurcating tree if the genomes within the hybrid lineage are considered independent but have population sizes and divergence times constrained to be identical (Jones et al. 2013). While subtle, the distinction between IM and MSCi approaches means that where IM models estimate dichotomous species trees while correcting for the distorting effects of migration, MSCi models altogether depart from strictly dichotomous trees in favour of a network with both speciation and reticulation edges (Bravo et al. 2019; Blair & Ané 2020; Cai & Ané 2021; Jiao et al. 2021). In this context, the standard MSC (i.e., the complete isolation model) can be considered a special case of the IM or MSCi in which the migration parameters in the species tree parameter vector have zero values.

The last decade has seen the development of multiple approximate-likelihood Bayesian implementations of the IM model, including the AIM extension to StarBeast2 (Müller et al. 2018, 2021), implementations in the IMa series in which the species tree topology is provided a priori (IMa2, Hey 2010; IMa2p, Sethuraman & Hey 2016) or inferred using a “hidden genealogy” approach (IMa3; Hey et al. 2018), MIST (Chung & Hey 2017), and DENIM (Jones 2019a), an extension of the standard MSC tree inference program DISSECT (Jones et al. 2015) with improvements implemented in STACEY (Jones 2017a). Analogous MSCi models include PhyloNet/MCMC-seq (Wen & Nakhleh 2018; Wen et al. 2018), SpeciesNetwork (Zhang et al. 2018) and a recent MSCi extension (Flouri et al. 2020) of BPP (Yang & Rannala 2010, 2014). The number of additional migration parameters estimated by IM and MSCi methods means that, like standard MSC methods (Bravo et al. 2019), tree or network inference for more than a handful of tips quickly becomes computationally demanding (Blair & Ané 2020; Cai & Ané 2021). In IMa2 (Hey 2010), MIST (Chung & Hey 2017) and the MSCi model implemented in BPP (Flouri et al. 2020), computational intensity can be mitigated by assuming that the species tree topology is known, so that sampling of alternative tree topologies is omitted from the MCMC. DENIM (Jones 2019a) mitigates computational load by integrating out the migration rates and population size parameters, and by ignoring non-parsimonious embeddings of gene trees in species trees. PhyloNet (Wen & Nakhleh 2018) and AIM (Müller et al. 2018, 2021) restrict migration to manageable levels by placing truncated and informative Poisson priors on the number of reticulation events and migration rates respectively (Blair & Ané 2020).

Even with simplifications to the tree space detailed above, using high throughput sequencing (HTS)-scale data sets of 200 or more loci with Bayesian IM and MSCi models is probably not computationally feasible (Blair & Ané 2020, Rannala et al. 2020). Particularly challenging is that thousands of loci may be necessary for reliable parameter estimation (Müller et al. 2018; Flouri et al. 2020; Rannala et al. 2020; Jiao et al. 2021), and each locus may need to be thousands of base-pairs (perhaps 10,000 bp) long for accurate joint inference of phylogeny and migration (Jones 2019a). Systematic evaluations of species tree inference under the standard MSC have demonstrated an exponential decline in tree estimation error and an exponential increase in computation time with an increase in the number of loci of fixed length (Heled & Drummond 2010; Ogilvie et al. 2016). Huang et al. (2020) recovered similar results for the estimation of divergence times, population sizes and introgression probabilities in the MSCi model as implemented in BPP (Flouri et al. 2020). In general, however, IM and MSCi models have been less well-studied than standard MSC tree inference methods (Degnan 2018; Blair & Ané 2020). Of particular concern is the lack of a systematic evaluation of IM-based inference assessing the effects of information content on tree estimation accuracy and computation intensity. Such an analysis would usefully inform experimental design or data subsampling regimes (Huang et al. 2020) in groups in which there is evidence that low to moderate levels of migration have contributed to overall gene tree discordance.

Here we address critical open questions in IM-based species tree inference, and more generally in contemporary molecular phylogenetics. We assess the impacts of information content on the computational efficiency and phylogenetic accuracy of IM and standard MSC models using simulated data sets comparable (in locus length and number) to those routinely obtained on medium- and long-read HTS platforms. Unlike previous work in which the rate of migration between branches is assumed to be uniform regardless of relatedness (Leaché et al. 2014a; Jones 2019a), we simulate sequences under the coalescent with migration that decays with time since the MRCA between contemporaneous branches. Our simulated pattern of migration rate decay reflects a general pattern of migration decline with time since the MRCA recovered in empirical studies of birds, mammals, fish, amphibians, and insects (Pinho & Hey 2010). We simulate coalescent sequences with continuous, low-level (< 0.01 migrants per generation), migration on an ultrametric ten-taxon tree (Leaché et al. 2014a; Jones 2019a) to assess tree inference under the IM model DENIM (Jones 2019a) and the standard MSC tree inference program StarBeast3 (Douglas et al. 2022). We quantify the impacts of molecular information content and migration on estimates of species tree topology and divergence times by DENIM, and compare computation time per unit ESS per CPU core as a measure of computational performance between DENIM and StarBeast3. We also assess the computational performance and phylogenetic accuracy of DENIM and StarBeast3 with an empirical data set assembled by Jebb et al. (2020) for six bat species.

## Materials & Methods

### Simulations

Gene trees were simulated along a rooted, ultrametric ten-taxon tree (Fig. 1a) following the MSC model (Rannala & Yang 2003). Simulated gene trees were used to simulate DNA sequences under a Jukes-Cantor substitution model (Jukes & Cantor 1969). Six simulation treatments were implemented, each representing a unique combination of locus number (30 and 60 loci) and locus length (600, 1000, and 10,000 base-pairs). Each treatment was replicated ten times. Sequences were simulated with the MCcoal program in BPP 3.4.0 (Rannala & Yang 2003; Yang & Rannala 2010) following the simulation approach of Leaché et al. (2014a) and Jones (2019a). Following Leaché et al. (2014a), speciation times (*τ*) and population sizes (θ) in replicate species trees were drawn from separate prior probability distributions (Fig. 1b-c) using values chosen to reflect those obtained in empirical studies (Leaché 2009, Castillo-Ramírez et al. 2010). Population sizes were sampled from an inverse gamma distribution with parameters α = 3, β = 0.03 and mean β/(1-α) = 0.015 (Fig. 1b), where population size is θ = 4*N_e_*μ, *N_e_* is the effective population size, and μ is the mutation rate per site per generation. Speciation times (*τ*), measured as the expected number of substitutions per site, were drawn from an exponential prior distribution with mean 1/λ = 0.02, such that the most recent species divergence in the tree is on average at *τ* = 0.02 and the root node height is on average at *τ* = 0.12 (Fig. 1a). To reduce variance among treatments, the ten replicate simulations conducted for each treatment assumed identical divergence times and population sizes but were independently seeded (Supplementary Material; DOI: 10.5061/dryad.cc2fqz6d1).

**FIGURE 1.**
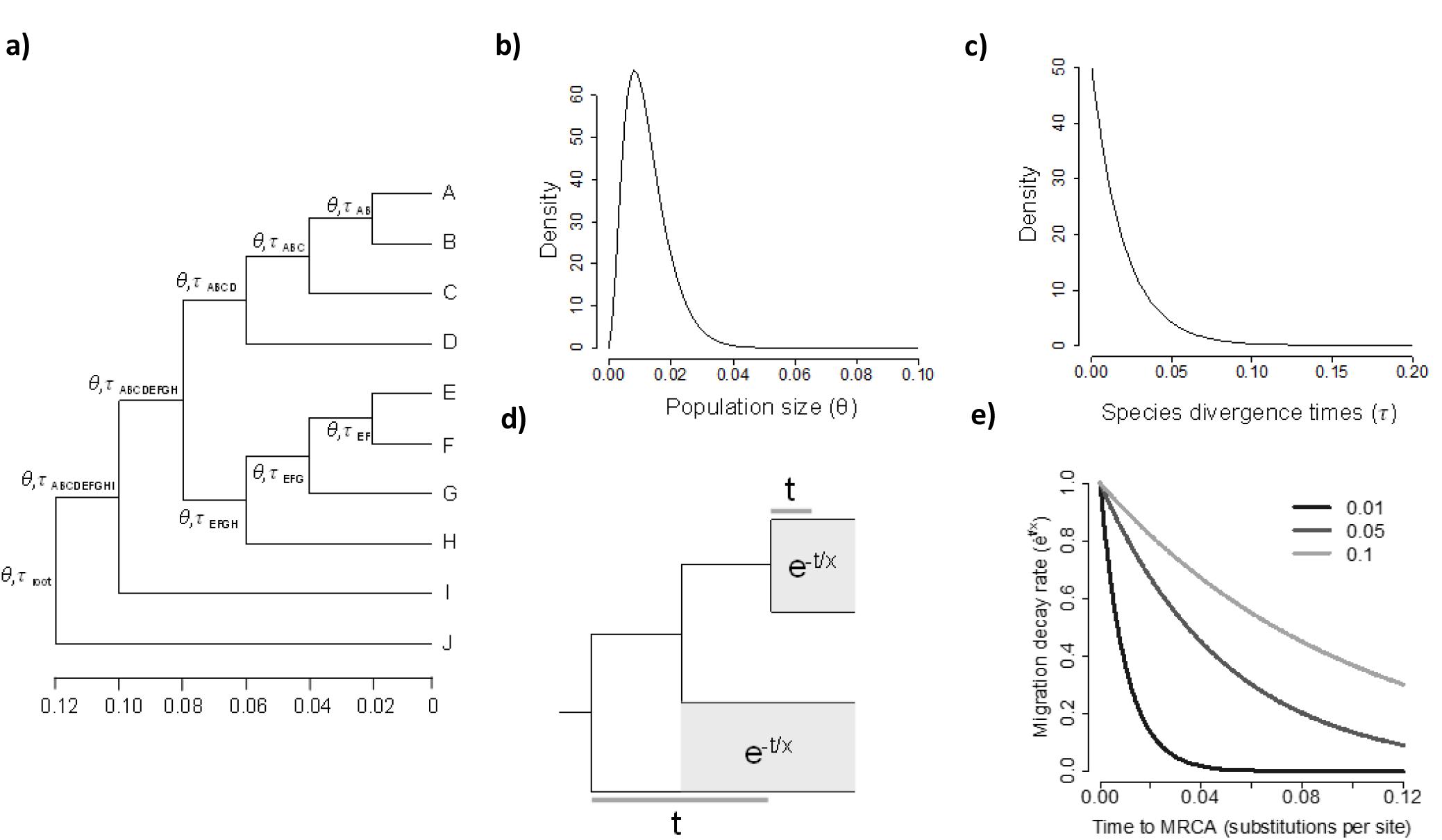
(a) Species tree topology along which coalescent sequences with migration were simulated using MCcoal, with time in expected substitutions per site. Ancestral nodes except the root are identified by the tip labels for which they represent the MRCA. (b) Inverse gamma distribution with mean 0.02 from which divergence times *τ* were sampled. (c) Exponential distribution with mean 0.015 from which population sizes as θ were sampled. (d) General model showing how migration rates between contemporaneous branches (branches connected by grey shading) are estimated according to rate *e^-t/x^*where *x* is the migration decay scale and *t* for each pair of branches is the time in expected substitutions per site between the MRCA and the midpoint of the time during which both branches exist. Grey areas show the time interval over which migrants are exchanged at a constant, migration decay scale-determined rate. Gene flow rates between pairs of branches eventually converge on zero so that there is an underlying species tree. (e) Migration decay rate *e^-x/t^*as a function of time *t* under alternative migration decay scale *x* values 0.01, 0.05 and 0.1.

**FIGURE 2.**
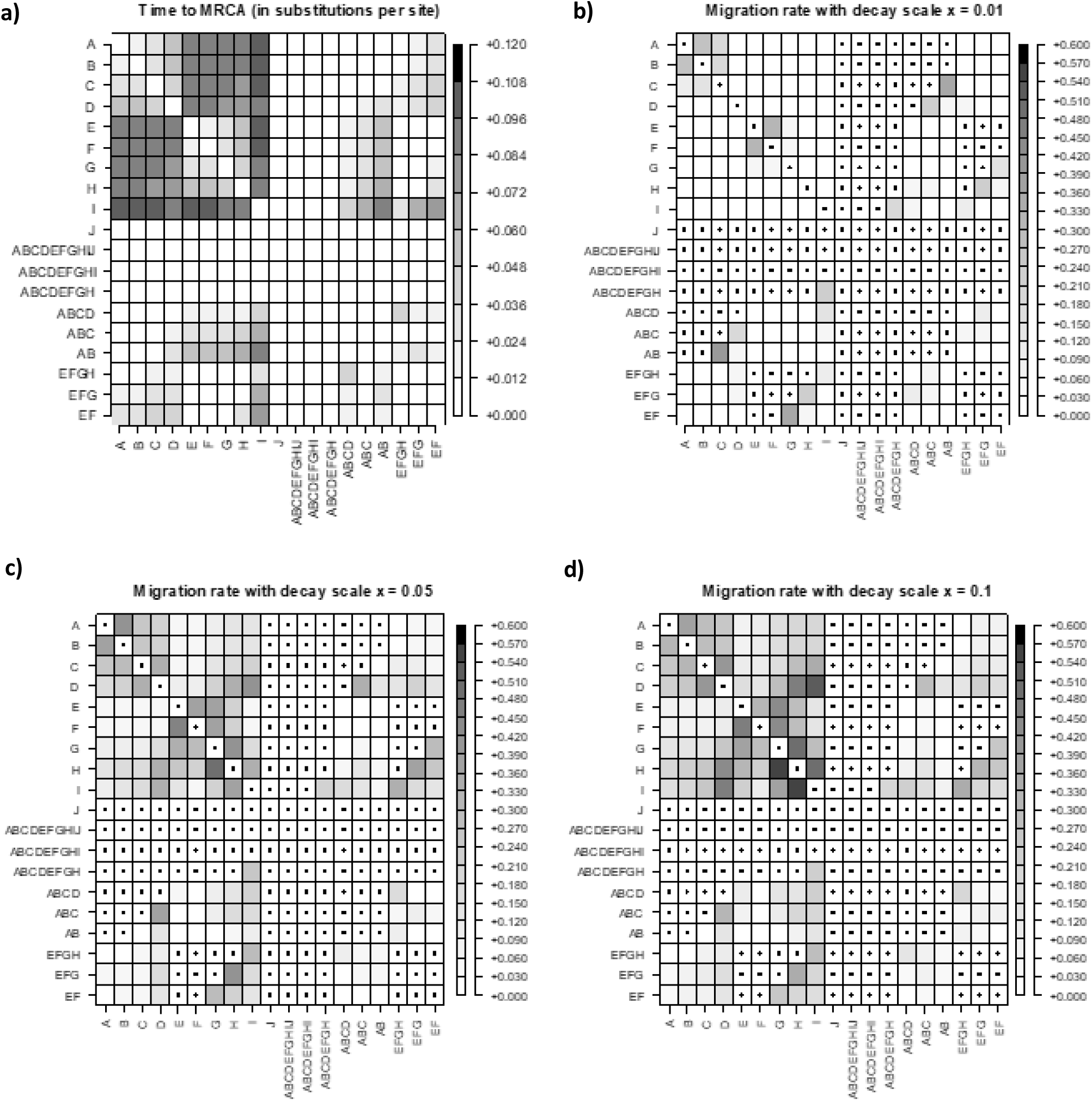
Migration rates between contemporaneous branches in simulated trees. (a) Time *t,* in expected substitutions per site, between the MRCA and the midpoint of the time during which contemporaneous migrant-exchanging branches exist and (b-d) mean simulated migration rates per generation between contemporaneous branches per unit branch length (in expected substitutions per site) for migration decay scale values x of 0.01, 0.05 and 0.1 respectively. Cells with centroids denote impossible (self, nested or non-contemporaneous) migrations.

We simulated continuous migration according to an n-island model with a migration matrix *M* = {*M_ij_*} where the scaled migration rate *M_ij_* = *N_j_m_ij_* is the expected number of migrants from lineage *i* to lineage *j* per generation, *m_ij_* is the proportion of immigrants in lineage *j* from lineage *i*, and *N_j_* is the effective population size of the receiving lineage *j* (Zhang et al. 2011; Jiao et al. 2021). Given that population size is related to time/branch length by θ = 4*N_e_*μ, the migration rate per sequence or lineage scaled by mutation (i.e., the number of migrants transferred along a branch of length 1 substitutions per site) is then 4*M_ij_*/θ*_j_* = *m_ij_*/μ (Jiao et al. 2021). Unlike the simulated sequences of Leaché et al. (2014a), the rate of migration between contemporaneous branches is not uniform across the tree, but instead decays exponentially as a function of time from the MRCA between migrant-exchanging lineages (Fig. 1d-e; Jones 2019b). This effect is achieved by multiplying a tree-wide migration rate, set to 0.01 migrants per sequence or lineage per generation (Leaché et al. 2014a; Jones 2019a), by *e^-t/x^*, where *t* is the time, in substitutions per site, between the MRCA of two branches and the midpoint of the time during which both branches exist, and *x* is the migration decay scale, the rate at which migration rates decrease with time since the MRCA between two lineages (Fig. 1d-e; Jones 2019a,b). In effect, the scaled rate of migrant exchange is constant over the time interval that two branches are contemporaneous (Fig. 1d), but decays in a stepped manner, with the intervening nodes as steps. To assess the influence of the migration decay scale value on tree inference, the ten replicate analyses per treatment were run with migration decay scale values of 0.01, 0.05 and 0.1. For sister species with an MRCA at node height 0.1 substitutions per site and constant population sizes θ of 0.015 (Fig. 1b), the expected number of migrants exchanged is therefore 1.80 x 10^−3^, 9.81 x 10^−2^ and 0.16 under migration decay scale values of 0.01, 0.05 and 0.1 respectively (Fig. 1e). As speciation times in replicate species trees are random samples from a prior distribution (Fig. 1c), the specific array of contemporaneous branches between which migration can occur varies in each replicate. We compiled a custom R script (R Core Team 2021; Supplementary Material) for computing branch length-dependent migration rate matrices. Four sequences were simulated per species, except for the outgroup species, for which only one sequence was simulated. Migration was simulated between all contemporaneous branches in the species tree, except for the outgroup species. Custom R scripts for the simulation data are in the Supplementary Material. We analysed additional simulated data sets with up to 2000 loci, but for these neither DENIM nor StarBeast3 achieved convergence after several weeks of computation (Supplementary Material).

### Empirical Data

We reconstructed phylogenetic relationships among the Laurasiatherian bat species *Molossus molossus* (Pallas, 1766)*, Myotis myotis* (Borkhausen, 1797), *Phyllostomus discolor* (Wagner, 1843), *Pipistrellus kuhlii* (Kuhl, 1817)*, Rhinolophus ferrumequinum* (Schreber, 1774), and *Rousettus aegyptiacus* (E. Geoffroy, 1810). Gene flow between non-sister species has been documented in *Myotis* (Morales & Carstens 2018) but, given that the six species under consideration diverged up to ca. 50 Ma (Moreno Santillán et al. 2021), we anticipated limited signal of intergeneric gene flow. Jebb et al. (2020) inferred a maximum likelihood species tree for the Laurasiatheria including these six bat species, based on a supermatrix of 12,931 orthologous protein-coding gene regions. We reanalysed this dataset with DENIM and StarBeast3 using a subset of 36 loci at least 10,000 bp long. We trimmed the sequences to 600, 1000, and 10,000 base-pairs to match the simulation sequence lengths. Each analysis was replicated twice. We analysed an additional empirical data set from Jebb et al. (2020) comprising 31 loci for seven primate species, but for these DENIM failed to achieve convergence after several weeks of computation (Supplementary Material).

### Phylogeny Inference with DENIM

We analysed the data sets described above using the IM program DENIM 1.0.0 (Jones 2019a) implemented in BEAST2 version 2.6.7 (Bouckaert et al. 2019) to infer a posterior distribution of species trees under the coalescent while accounting for low to moderate levels of migration (< 0.01 migrants per generation) among contemporaneous branches. As the species tree is of primary interest, DENIM integrates out population size and migration parameters, but estimates of migration within individual loci are retained and can be retrieved by postprocessing. The tree space explored by the MCMC is further reduced with an approximation wherein non-parsimonious gene tree embeddings in species trees are ignored so that the number of expected migration events per locus cannot exceed the number of coalescent events in the tree. This approximation holds provided migration between branches remain below one migrant per ten to 100 generations (Jones 2017b-c, 2019a), as in our simulations. Where migration rates exceed this range, the ignored non-parsimonious embeddings of gene trees in the species tree become increasingly likely, and the approximation breaks down.

DENIM XML files were prepared in BEAUTi 2.6.7 (Bouckaert et al. 2019). XML files for larger analyses comprising hundreds of loci were manually assembled in R using a custom script (Supplementary Material). Site models were linked among loci. Clock models were strict and unlinked with the first clock rate = 1 and rates for the remainder of loci estimated. A birth-death model was used to model the species tree. The nucleotide substitution model was set to JC69 to match the simulated model. The shape (α) and scale (β) of the migration rate prior (GammaComponent.1) were set to 3.0 and 0.02 to achieve a mean prior β/(1-α) = 0.01. Relatedness Factor was set to −1 so that migration between contemporaneous lineages is unrelated to the number of rootward branches separating taxa. Migration decay scale was set to 0.01, 0.05 and 0.1 according to the values under which sequences were simulated. For the bat sequence data, migration decay scale was set to 0.01. Species limits were fixed by setting Collapse Weight to 0 and changing the default prior to - 0.5. Uniform priors were replaced with lognormal priors for the relative clock rates: lognormal(0,1), growth rate for the species tree: lognormal(5, 2), and popPriorScale: lognormal(−7, 2; Jones 2019a, 2019b). The prior for relativeDeathRate was uninformative with a beta distribution β(1, 1). Each DENIM analysis was run for 300 x 10^6^ generations, storing every 100 000^th^ tree with 10% discarded burnin. Where effective sample sizes (ESS) for the posterior were below 200, the MCMC was extended by 300 x 10^6^ generations until an ESS of at least 200 was achieved. To accelerate computation, gene tree loggers were removed from all XML files. Each analysis was run on a single Intel Xeon Gold 6130 processor with 15G RAM on the *Tetralith* cluster at the National Supercomputer Centre (www.nsc.liu.se) under projects SNIC 2021/5-247, SNIC 2022/5-276 and NAISS 2023/5-403.

### Phylogeny Inference with StarBeast3

The standard MSC model, which infers the species trees without accounting for migration, was implemented for the data sets described above using the BEAST2 program StarBeast3 1.0.4 (Douglas et al. 2022). StarBeast3 XML files were prepared in BEAUTi 2.6.7 (Bouckaert et al. 2019) and XML files for larger analyses were assembled using a custom R script (Supplementary Material) with settings chosen to optimize similarity with the DENIM analyses. Simulated molecular data, site models, clock models, and nucleotide substitution models were identical to those in the DENIM analyses. A birth-death model was used for the species tree prior with uninformative priors for ExtinctionFraction: β(1, 1) and netDiversificationRate: Lognormal(5, 2), and the prior for mean effective population size lognormally distributed: Lognormal(−7, 2). Each StarBeast3 analysis was run for 300 x 10^6^ generations, storing every 100 000^th^ tree with 10% discarded as burnin. Where effective sample sizes (ESS) for the posterior were below 200, the MCMC was extended by 300 x 10^6^ generations until an ESS of at least 200 was achieved. For each StarBeast3 run, four kernels were used. To accelerate computation, gene tree loggers were removed from all XML files. Each analysis was parallelized across four Intel Xeon Gold 6130 processors with 15Gb RAM on the cluster *Tetralith*.

### Evaluation Measures

Several measures were used to assess phylogenetic accuracy. Topology estimation was assessed with the posterior probability of the true (i.e., simulated) topology. Divergence time estimation was evaluated by comparing the difference in the true (i.e., simulated) and estimated heights of node *τ_ABCDEFGHI_*, the MRCA of all migrant-exchanging branches. We also jointly assessed topology and branch length estimation accuracy using the branch score distances of Kuhner and Felsenstein (1994) adapted for rooted trees (Heled & Drummond 2010, Equation [9]). For all evaluation measures, we use the full posterior sample of trees after removal of between 10% and 80% burn-in (Supplementary Material). The 95% credible set of species trees was summarized for each posterior sample with the BEAST 1.10.4 program TreeLogAnalyser (Suchard et al. 2018). Branch scores were computed with the R package *phangorn* 2.10.0 (Schliep 2011). Computation time was measured as time per unit posterior ESS per core. MCMC trace logs were summarised in Tracer 1.7.2 (Rambaut et al. 2018). The tree-wide simulated number of migration events was calculated as the sum of 4*M_ij_*/θ*_j_* for all migrant-exchanging pairs standardised by the actual branch length (in expected substitutions per site) two branches were contemporaneous. Results from DENIM and StarBeast3 were analysed jointly with factorial ANOVAs in the R base package *stats* (Supplementary Material) with tree inference method, locus length and number, and migration decay scale as categorical predictor variables. Scripts for computing and displaying the evaluation measures are in the Supplementary Material.

## Results

### Simulations

Of the total 360 replicates we analysed, DENIM required an average of 3.16 x 10^8^ generations to achieve a posterior ESS of 200, while StarBeast3 required an average of 3 x 10^8^ generations (Supplementary Material). All analyses were continued until a posterior ESS of 200 was reached, with StarBeast3 converging at 20% the rate of DENIM per hour per core on average (Fig. 3a-b; F_(1)_ = 160.92, p < 0.0001). Increasing the number of loci from 30 to 60 significantly slows convergence under both DENIM and StarBeast3 (F_(1)_ = 119.94, p < 0.0001), but neither sequence length (F_(2)_ = 0.47, p = 0.627) nor tree-wide migration rate (F_(2)_ = 2.97, p = 0.053) have a significant effect on the rate of model convergence (Supplementary Material).

**FIGURE 3.**
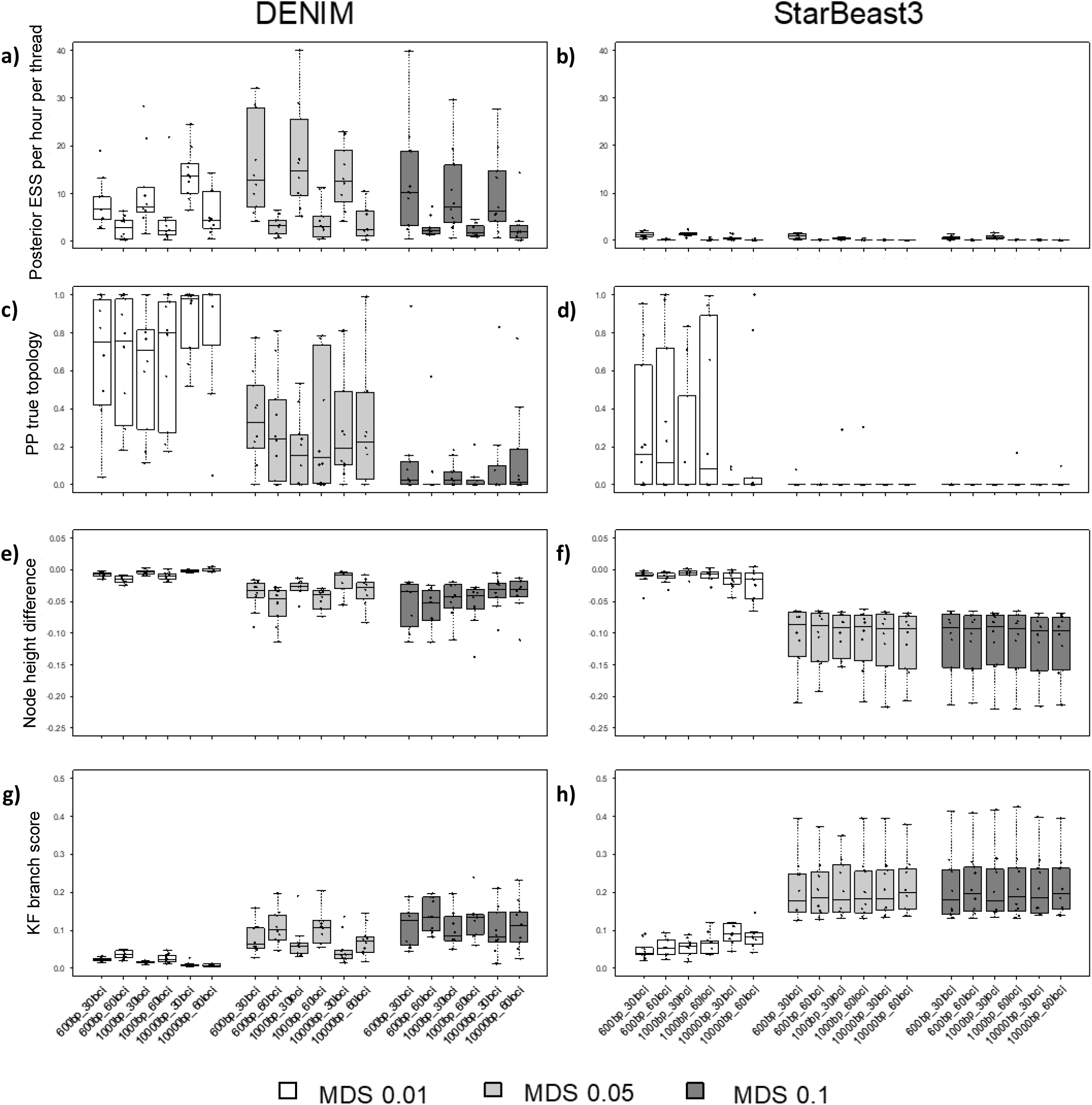
Evaluation measures for DENIM and StarBeast3 with simulated sequences for ten replicates. (a-b) Posterior density ESS per hour per core after burn-in removal. (c-d) Posterior probability of the true (i.e., simulated) topology in the posterior sample of trees. (e-f) Difference between the simulated and estimated node height *τ*_ABCDEFGHI_ in expected substitutions per site. (g-h) Kuhner and Felsenstein (1994) branch score distances between the true (i.e., simulated) and estimated trees. Legend denotes migration simulation at migration decay scale *x* values 0.01, 0.05 and 0.1. Boxplots span the interquartile range of values in the response variable. The middle 50 % of the data lie within the box, with the line in the box representing the median. Minimum and maximum values are depicted with whiskers extending from the box.

Figures 3c and 3d show the posterior probability of the true species topology in the posterior sample of trees. Under both DENIM and StarBeast3, the true topology is recovered with higher posterior probability under lower tree-wide migration rates (F_(2)_ = 104.94, p < 0.0001). DENIM retrieves the correct topology with higher posterior probability than StarBeast3 overall (F_(1)_ = 126.43, p < 0.0001), with StarBeast3 retrieving the true topology with posterior probabilities near zero even under relatively low migration rates of < 0.001 migrants per generation. Neither sequence length (F_(2)_ = 0.35, p = 0.703) nor the number of loci (F_(1)_ = 0.61, p = 0.436) has a significant effect on topology estimation (Supplementary Material).

Figures 3e and 3f show the difference between the true (simulated) and estimated heights of node *τ_ABCDEFGHI_*. DENIM and StarBeast3 underestimate the height of node *τ_ABCDEFGHI_* at all levels of migration, with the underestimation worsening under slower migration decay (F_(2)_ = 177.18, p < 0.0001). The underestimation of divergence times among migrant-exchanging branches is significantly worse in StarBeast3 (F_(1)_ = 192.06, p < 0.0001), where the mean height of node *τ_ABCDEFGHI_* is 0.0417 (sd = 0.055) substitutions per site. By comparison, the mean height of node *τ_ABCDEFGHI_* is 0.0882 (sd = 0.0453) substitutions per site under DENIM, nearer the mean true (i.e., simulated) height 0.119 (sd = 0.052) substitutions per site.

Figures 3g and 3h show the Kuhner & Felsenstein (1994) branch scores capturing topology and branch length estimation accuracy. Branch scores for both DENIM and StarBeast3 worsen under elevated levels of migration (F_(2)_ = 163.21, p < 0.0001). Consistent with trends in topology and divergence time estimation, DENIM outperforms StarBeast3 (F_(1)_ = 235.579, p < 0.0001) under all levels of migration.

Migration rates are integrated out by DENIM, but the total number of migration events over the tree can be retrieved by postprocessing (Jones 2019a). Figure 4 shows the mean number of migration events per locus inferred over the entire posterior sample of trees. The number of migration events detected by DENIM increases significantly where more migration is simulated (F_(2)_ = 467.86, p < 0.0001), demonstrating DENIM’s ability to detect even subtle changes in migration rate. Although the simulated and inferred number of migration events do not correlate for any of the 18 simulation conditions, the inferred number of migration instances across migration levels falls within the range of simulated values under all treatments (Fig. 4, Supplementary Fig. S1). While the mean tree-wide number of simulated migration events was 0.53 (sd = 0.57), 4.71 (sd = 4.77) and 7.71 (sd = 7.39) under MDS values of 0.01, 0.05 and 0.1 respectively, the inferred number of migration events was 1.14 (sd = 1.26), 4.83 (sd = 1.66) and 6.89 (sd = 1.70). All other conditions being equal, increasing the number of loci from 30 to 60 increases the number of inferred migration events 1.44 times (F_(1)_ = 37.21, p < 0.0001).

**FIGURE 4.**
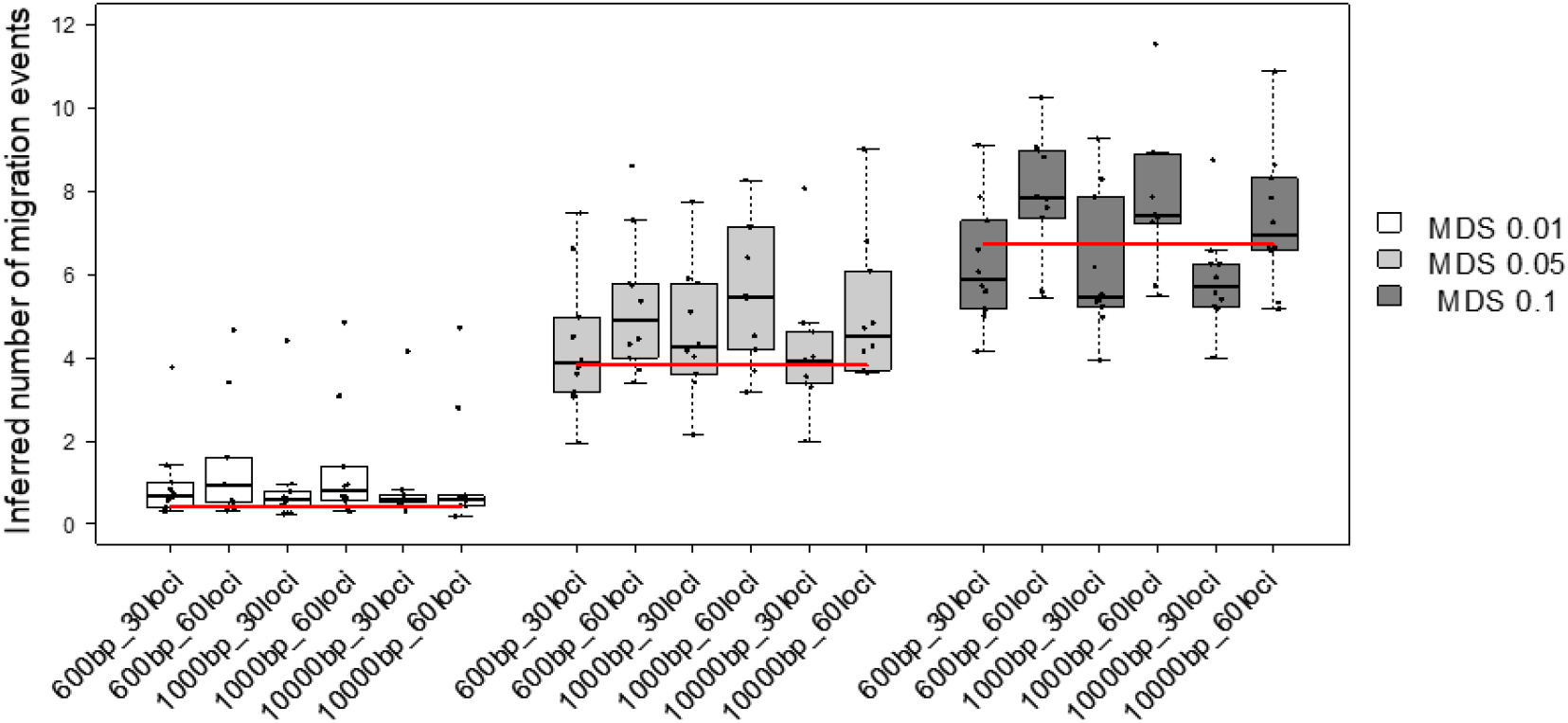
Number of inferred migration events per-locus by DENIM with simulated sequences for ten replicates. Red horizontal line represents the median simulated number of migration events per locus. Legend denotes migration simulation at migration decay scale *x* values 0.01, 0.05 and 0.1. See Figure 3 caption for description of boxplots.

### Empirical Data

Our DENIM and StarBeast3 analyses of a 36-locus subset of the Laurasiatherian bat sequence data agrees topologically with the maximum likelihood concatenated phylogeny estimated by Jebb et al. (2020; Supplementary Material). Table 1 shows that DENIM converges faster than StarBeast3 per core, especially with 10,000 base-pair long loci.

**TABLE 1.** Evaluation measures for the 36-locus bat data set under DENIM and StarBeast3.

|  |  | DENIM |  | StarBeast3 |  |
| --- | --- | --- | --- | --- | --- |
|  | Sequence length (bp) | Replicate 1 | Replicate 2 | Replicate 1 | Replicate 2 |
| Posterior ESS per hour per core | 600 | 38.98 | 49.30 | 11.82 | 11.67 |
|  | 1000 | 31.33 | 33.13 | 10.72 | 10.86 |
|  | 10,000 | 591.58 | 257.63 | 0.38 | 0.42 |
| Posterior probability<sup>a</sup> | 600 | 0.89 | 0.87 | 0.89 | 0.88 |
|  | 1000 | 0.98 | 0.99 | 0.96 | 0.95 |
|  | 10,000 | 1 | 1 | 1 | 1 |
| Estimate height of the root node in the MCC tree (subst. per site) | 600 | 0.0454 | 0.0454 | 0.0466 | 0.0466 |
|  | 1000 | 0.0579 | 0.0578 | 0.0569 | 0.0571 |
|  | 10,000 | 0.1109 | 0.1050 | 0.0858 | 0.0849 |

<sup>a</sup> Fraction of trees in the posterior sample matching the Jebb et al. (2020) ML tree topology.

Consistent with the simulated results, StarBeast3 recovers shallower root node heights than DENIM, but only clearly so with 10,000 base-pair loci. The posterior probabilities of the tree topology recovered by Jebb et al. (2020) are similar for DENIM and StarBeast3, with both methods recovering higher posterior support with longer sequences.

## Discussion

Our simulation work delivers three insights relevant to contemporary molecular phylogenetics. Firstly, the IM method DENIM outperforms the standard MSC method StarBeast3 in terms of phylogenetic accuracy and per core computational performance in the presence of low-level migration (Fig. 3). Jones (2019a) recovered similar results in his implementation of the standard MSC approach *BEAST (Heled & Drummond 2010) under migration, with *BEAST almost never recovering the true topology under small amounts of non-sister migration. The misestimation of branch lengths under standard MSC methods in the presence of migration is also well-documented (Leaché et al. 2014a; Long & Kubatko 2018; Müller et al. 2018; Wen and Nakhleh 2018; Elworth et al. 2019, Jiao et al. 2020, Pang & Zhang 2022). As for computational performance, DENIM converges up to an order of magnitude faster than StarBeast3, a surprising result given that DENIM includes additional migration parameters that are absent from StarBeast3. In the absence of migration and with the same number of threads, however, StarBeast3 appears to achieve stationarity faster (Douglas et al. 2022), which suggests that its relatively poor performance is a consequence of model violation. We note that StarBeast3 was designed to optimize computational efficiency by multi-threading gene tree estimation, where our largest data set comprised only 60 loci run on four cores. We also note that StarBeast3 estimates effective population size parameters during the MCMC sampling process, while DENIM integrates out these parameters completely.

The second insight relates to the rate at which migrant exchange decays as a function of time from the MRCA between lineages. Rapidly decaying migration is associated with better topology and branch length estimates in both DENIM and StarBeast3, with DENIM outperforming StarBeast3 under all migration decay scenarios (Fig. 3c-h). The deteriorating performance of StarBeast3 under elevated migration can possibly be attributed to more intensive model violation under gradual migration decay. By contrast, DENIM’s deteriorating performance (Fig. 3c-h) under gradually decaying migration is potentially due to a breakdown in its parsimony approximation around unlikely embeddings of gene trees in the species tree, but our results indicate that the estimated number of migration events per locus remains well below the upper limit (viz. the number of coalescent events) even under the highest levels of migration simulated (Fig. 4; Supplementary Material). At any rate, the parsimony approximation around unlikely embeddings of gene trees in species trees implemented in DENIM performs well where migration decay is rapid.

The third insight relates to phylogeny estimation under varying information content. Doubling locus number from 30 to 60 significantly slows MCMC mixing without significantly improving topology or branch length estimates. By contrast, increasing sequence length from 600 to 10,000 base-pairs does not significantly reduce convergence rate, but results in subtle improvements in the accuracy of branch length and topology estimates. Consistent with Jones’ (2019a) findings, our results argue for the use of longer, rather than more, loci, and although longer sequences are likelier to include points of recombination, previous work suggests MSC-based species tree estimation is robust to realistic levels of recombination (Lanier & Knowles 2012, Lohse & Frantz 2014, Zhu et al. 2022). A more immediate challenge for molecular phylogeneticists, perhaps, is the cost effectiveness of sequencing and assembling long sequences for many accessions. While short-read sequencing is relatively cost-effective, using short reads to resolve repetitive and complex regions remains challenging. Sequencing platforms by PacBio, ONT, Element Biosciences, Illumina and MGI overcome these challenges by sequencing longer reads (15kbp–4Mbp, De Coster et al. 2021, Marx 2023). For future applications of the IM model, targeted enrichment followed by long-read sequencing may represent a viable solution for scaling up the assembly of long sequences (De Coster et al. 2021), perhaps even for degraded museum specimens (Quatela et al. in press).

Simulation studies assessing the performance of IM and MSC inference methods have hitherto assumed that the migration rate between a pair of lineages is unrelated to the genetic distance between them, even as the time to the MRCA approaches 0.1 substitutions per site (Leaché et al. 2014a; Jones 2019a). Empirically, this implies equal rates of migrant exchange among all water birds (Hackett et al. 2008), among cetaceans (Liu et al. 2010), among humans, great apes, and other simians (Perelman et al. 2011), among spiny rats (Upham et al. 2013), and among fungus-growing ants (Ward et al. 2015). Here we introduce a more biologically plausible approach to IM sequence simulation in which the probability of migrant exchange is higher between close relatives. This corresponds more closely to observed patterns of interspecific gene flow in amphibians (Malone & Fontenot 2008), birds (Tubaro & Lijtmaer 2002, Arrieta et al. 2013), flowering plants (Moyle et al. 2004, Scopece et al. 2007), fungi (Giraud & Gourbière 2012), and insects (Orr 2005, Sánchez-Guillén et al. 2014, Turissini et al. 2018; see Pinho & Hey 2010). We further advance the biological realism of our simulated sequences by removing individual-, clade-, and allele-specific restrictions on migrant exchange (e.g., Leaché et al. 2014a, Long & Kubatko 2018, Jones 2019a), and instead simulate sequences with migration between all contemporaneous branches.

The last decade has seen the development of two major extensions of the standard MSC: the MSC with delimitation of Wright-Fisher species, and the MSC with cross-species gene flow, the latter including continuous-time migration modelled under IM, and episodic migration modelled under the MSCi. MSC-based approaches for species discovery or validation have been widely applied, and often inform taxonomic revisions (e.g., Leaché & Fujita 2010; Leaché et al. 2014b; Toprak et al. 2016; Noguerales et al. 2018). By contrast, IM and MSCi methods have remained relatively underutilized, probably due to their more intensive parameterization and perceived computational inefficiency (although see Jackson et al. 2017). In a recent review, Rannala et al. (2020) suggested that reliable migration parameter estimates under IM and MSCi methods may require large data sets comprising thousands of loci, and that the use of realistically sized data sets comprising ca. 200 loci is probably computationally unfeasible. Our work demonstrates that DENIM can deliver accurate phylogeny estimates in the presence of both ILS and empirically realistic patterns of migration using as few as 30 loci. Moreover, under all sequence sampling strategies implemented here and under rapidly and gradually decaying patterns of tree-wide migration, DENIM computationally outperforms StarBeast3. While StarBeast3 may be able to achieve convergence faster than DENIM where the number of processors approaches the number of gene trees (see Douglas et al. 2022), further parallelizing StarBeast3 cannot address the systematic misestimation of topology and underestimation of speciation times where signal of migration is present (Leaché et al. 2014a; Long & Kubatko 2018; Müller et al. 2018; Wen and Nakhleh 2018; Elworth et al. 2019, Jiao et al. 2020, Pang & Zhang 2022). Our work provides analytical support for the broader application of IM methods where signature of postspeciation gene flow is detectable, not only as a means of mitigating migration-induced distortions in the species tree, but of accelerating computation.

## Supplementary Material

Data available from the Dryad Digital Repository: https://datadryad.org/stash/share/BPNSSnghL3Igx6ynL0Rq-srxyqjreiqKfjB2nlmjOH4

## Acknowledgements

The authors are thankful to Graham Jones for assistance with DENIM and for detailed comments on this manuscript. Adam Leaché and Ziheng Yang are thanked for assistance with MCcoal. Scott Edwards and Bruce Rannala are thanked for valuable feedback. Computation was enabled by resources provided by the National Academic Infrastructure for Supercomputing in Sweden (NAISS) and the Swedish National Infrastructure for Computing (SNIC) on cluster Tetralith partially funded by the Swedish Research Council through grant agreements 2022-06725 and 2018-05973. Mats Kronberg at the National Supercomputer Centre is thanked for assistance with technical aspects of the work on Tetralith.

